# Visual looming and receding networks in awake marmosets investigated with fMRI

**DOI:** 10.1101/749309

**Authors:** Justine C. Cléry, David J. Schaeffer, Yuki Hori, Kyle M. Gilbert, Lauren K. Hayrynen, Joseph S. Gati, Ravi S. Menon, Stefan Everling

## Abstract

An object that is looming toward a subject or receding away contains important information for determining if this object is dangerous, beneficial or harmless to them. This information (motion, direction, identity, time-to-collision, size, velocity) is analyzed by the brain in order to execute the appropriate behavioral responses depending on the context: fleeing, freezing, grasping, eating, exploring. In the current study, we performed ultra-high-field functional MRI (fMRI) in awake marmosets to explore the patterns of brain activation elicited by visual stimuli looming toward or receding away from the monkey. We found that looming and receding visual stimuli both activate a large cortical network in frontal, parietal, temporal and occipital cortex in areas involved in the analysis of motion, shape, identity and features of the objects. Looming stimuli strongly activated a network composed of the pulvinar, superior colliculus, prefrontal cortex and temporal cortical areas. This may underlie the existence of an alert network that processes the visual stimuli looming toward their peripersonal space by extracting the crucial information brought by the stimulus and evaluating its potential consequences to the observer. We hypothesize that this network is involved in the planning of protective behaviors (e.g. fleeing or freezing) and in emotional reaction (e.g. anxiety, fear). These findings support the view that this network is preserved through evolution and that the marmoset is a viable model to study visual and multisensory processes by using fMRI to guide further invasive recordings and/or pharmacological manipulations.

**Significant statement:** An object that is looming toward a subject or receding away contains important information for determining if this object is dangerous, beneficial or harmless to them. Here, we identified the functional network in non-human primates that was activated by visual stimuli looming toward or away from the animals using ultra-high-field functional magnetic resonance imaging. Our findings show that large cortical activations are elicited by both looming and receding visual conditions. However, some activations were specific to the looming condition, suggesting that the integration of cues in the looming direction rely on strong connections between cortical and subcortical areas, which allows primates to react properly for protecting themselves against a potential threat.

## Introduction

The primate’s ability to perceive motion such as looming (e.g. a predator) or receding stimuli (e.g. a prey) is essential for adapting and adjusting their behavior to the context (e.g. fleeing, freezing or hunting). Looming stimuli are characterized by the expansion of a closed contour in the field of view, and alternatively, receding stimuli are characterized by contraction of a closed contour (Schiff et al., 1962). Franconeri and Simons (2003) proposed a behavioral-urgency hypothesis: the processing of stimuli are prioritized for those that signal an event that could require immediate action such as adaptive behavior or defensive responses (Náñez, 1988; Náñez and Yonas, 1994; Shirai et al., 2004). In both infants and adult Old World macaque monkeys and humans, looming stimuli trigger stereotypical avoidance responses such as leaping, springing or ducking, whereas receding stimuli elicit exploratory behaviors (Schiff et al., 1962; Ball and Tronick, 1971). Such a behaviour requires integrating and analyzing the spatiotemporal components, the depth cues, the relative distance and the direction of approach in a short period of time.

Over the past fifteen years, behavioral studies have shown that the predictive mechanisms underlying the approach of stimuli toward the observer involve multisensory processes such as, in most cases, the stimulus predicting a collision leading to a tactile impact on the body. Behavioral orienting indices are increased by auditory-visual stimuli (Maier et al., 2004; Cappe et al., 2009). For example, auditory or visual looming stimuli both enhance tactile processing at the predicted time of impact and at the expected location of impact by increasing tactile sensitivity (Cléry et al., 2015a) and by shortening reaction times (Canzoneri et al., 2012; Kandula et al., 2015). Recently, an fMRI study performed in macaque monkeys (Cléry et al., 2017) highlighted the existence of a core cortical network of prefrontal, premotor, parietal and visual areas involved in the processing of visual looming stimuli predicting an impact into the face. This network has been suggested to be involved in approaching behavior (Rizzolatti et al., 1997) and in the representation of a peripersonal space for serving defense and protective behaviors (Graziano and Cooke, 2006; Cléry et al., 2015b), as confirmed by the partial overlap of this network with the network encoding peripersonal space (Cléry et al., 2018). This mechanism seems conserved across macaques and humans suggesting that it is a conserved network as their common ancestor lived about 25 million years ago (Miller et al., 2016). Further invasive explorations using high density and laminar recordings and manipulations are needed for a better understanding of the neuronal mechanism in this network. This type of studies is difficult or even impossible in macaque monkeys as many of the frontal and parietal regions are located deep within sulci (arcuate sulcus and intraparietal sulcus).

With a largely lissencephalic cortex, the New-World common marmoset (*Callithrix jacchus*) may be a potentially alternative primate model for studying the neural processes occurring in frontal and parietal areas during looming and receding visual stimulation. A prerequisite for invasive recording studies is the identification of the brain areas involved in the processing of looming and receding stimuli in marmosets. Here we took advantage of the small size of these primates and used a small-bore ultra-high-field 9.4T MRI scanner to explore the whole-brain level fMRI activations elicited by visual stimuli looming toward or receding away from awake marmosets.

## Methods

All procedures were in compliance with the Canadian Council of Animal Care policy. All protocols used in this experiment were approved by the Animal Care Committee of the University of Western Ontario Council on Animal Care.

### Subjects and experimental setup

Three male common marmosets (*Callithrix jacchus*) were involved in this study. Their age and weight at the time of the experiments was 36 months and 380 g (M1), 20 months and 360g (M2) and 19 months and 245g (M3), respectively. The animals were prepared for awake fMRI experiments by implanting an MRI-compatible head restraint/recording chamber (for details, see Johnston et al., 2018). Moreover, prior to each imaging session, the head chamber was filled with a water-based lubricant gel (MUKO SM321N, Canadian Custom Packaging Company, Toronto, Ontario Canada) to reduce the magnetic-susceptibility image artifacts created by the skull-attached chamber (for details, see Schaeffer et al., 2019). However, it should be noted that the coverage of visual cortex is not optimal as the temporal signal-to-noise ratio dropped in this part of the brain (see Fig. 4f in Schaeffer et al., 2019).

During the scanning sessions, the animal sat in the sphinx position in an animal holder consisting of an MRI-compatible restraint system (Schaeffer et al., 2019) with an MRI-compatible camera (Model 12M-i, 60-Hz sampling rate, MRC Systems GmbH, Heidelberg, Germany) and a reward tube at the front. The animals were first restrained using neck and tail plates, then the head was restrained by the fixation of the head chamber to the five-channel receive coil. The animal was monitored during scanning by a veterinary technician using the MRI-compatible camera. The animals were rewarded with diluted sweet banana milk dispended by an injection pump (New Era Pump Systems Inc., Model NE-510, Farmingdale, NY USA) after each block to keep them motivated and alert throughout data acquisition. In the MR scanner, the animal faced a translucent plastic screen placed at the front of the bore (119 cm from the animal head) where visual stimuli were displayed via back-projection through a mirror with a SONY VPL-FE4 projector. The maximum angle-of-view from the center to the side of the screen was 7°. The task, the reward delivery and the visual stimulations were synchronized from the MRI trigger with a custom-written program running on a Raspberry Pi (Model 3 B+, Raspberry Pi Foundation, Cambridge, UK). Monkeys were habituated to the MRI scanner setup in a mock scan environment for about two weeks (Schaeffer et al., 2019).

### Task and stimuli

A dot was placed in the center of a simulated 3D environment with visual depth cues and was present throughout each run (Fig.1A). The 3D environment and stimuli were all constructed with the Blender software (http://www.blender.org/). Visual stimuli consisted of dynamic 3D geometrically shaped stimuli rotating upon themselves. There were seven different shapes: a cube, a cylinder, a ball, a simple icosphere, a complex icosphere, a torus and a cone pointing toward the marmoset (Figure 1A). We used different stimuli to maintain the animal arousal level throughout acquisition. Visual stimuli loomed toward the animals or receded away from them (at 17.33 cm/s). Looming and receding conditions were presented in the same runs but in separate blocks of eleven stimuli each. The marmosets were not required to fixate on the dot or track the stimuli during the task.

**Figure 1:**
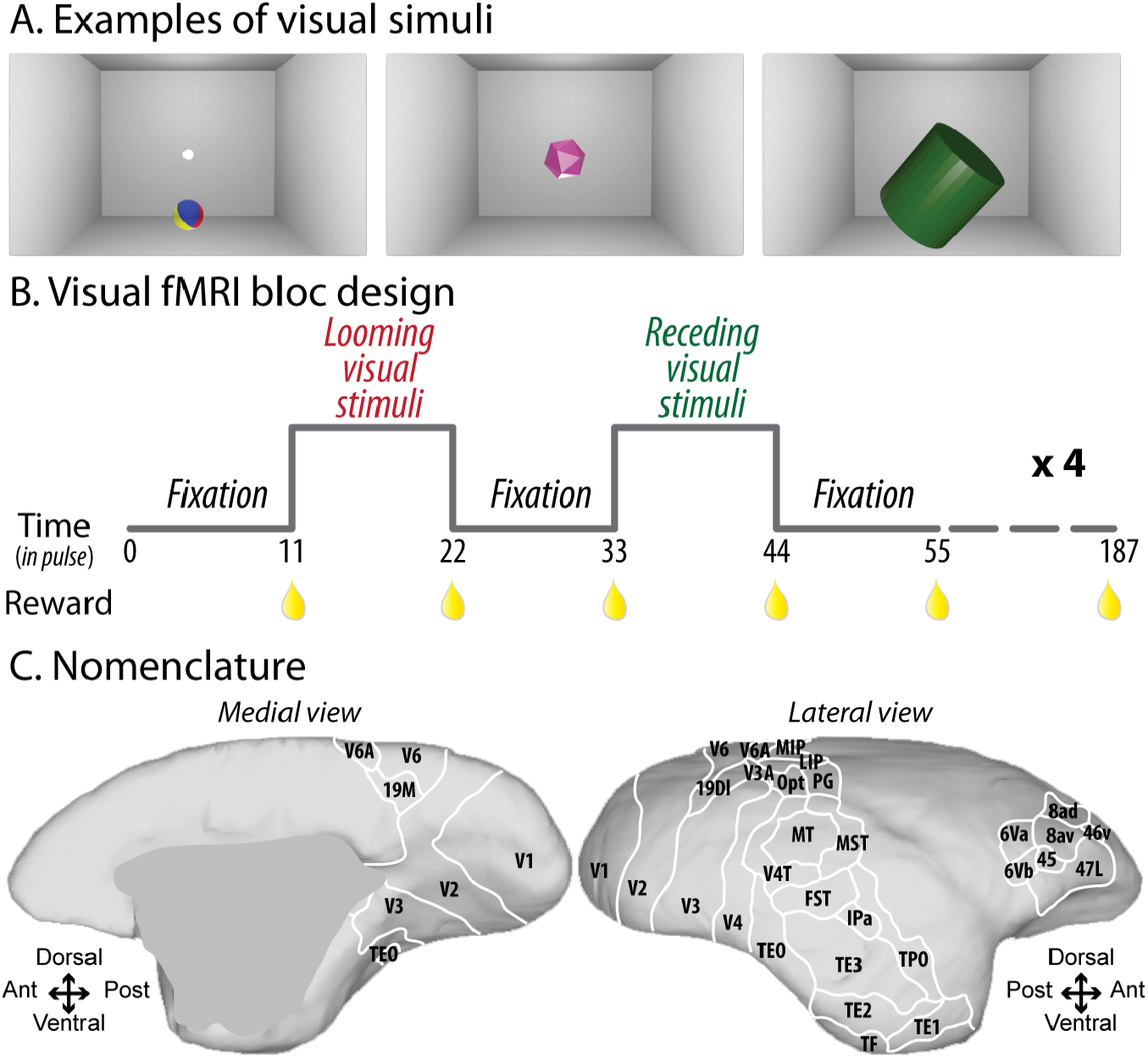
Experimental fMRI protocol. A) Examples of visual stimuli displayed during the looming or receding condition. In total, seven different objects were presented: a cube, a cylinder, a ball, a simple icosphere, a complex icosphere, a torus and a cone pointing toward the marmoset. B) Visual fMRI block design. It consisted of a succession of blocks in which the object was moving toward the animal and blocks in which the object was moving away from the animal, separated by fixation blocks. This pattern was repeated four times: each block lasted 11 pulses and a reward drop (banana milk) was provided to the animal after each block. C) The nomenclature of the cortical regions was based on the histology-based atlas of Paxinos and colleagues (2011).

Functional time series (runs) were organized as follows (Fig. 1B): an 11-volume block of no stimuli was followed by an 11-volume block of visual stimuli looming toward the monkey, an 11-volume block of no stimuli, an 11-volume block of visual stimuli receding away from the monkey and an 11-volume block of no stimuli (baseline). A given sequence, repeated four times, resulted in a 187-volume run with 17 blocks.

### Scanning

We performed data acquisition using a 9.4-T, 31-cm horizontal-bore magnet (Varian/Agilent, Yarnton, UK) and Bruker BioSpec Avance III console with the software package Paravision-6 (Bruker BioSpin Corp, Billerica, MA) at the Centre for Functional and Metabolic Mapping at the University of Western Ontario. We used an in-house, custom-built integrated receive coil with five channels (Schaeffer et al 2019) paired with a custom-built high-performance 15-cm-diameter gradient coil with 400-mT/m maximum gradient strength (xMR, London, CAN; Peterson et al., 2018). We used an in-house quadrature birdcage coil (12-cm inner diameter) for the transmit coil.

For each animal, a T2-weighted structural image was acquired, to allow an anatomical registration, with the following parameters: repetition time (TR) = 5500ms; echo time (TE) = 53 ms; field of view (FOV)= 51.2 × 51.2 mm; voxel size = 0.133 × 0.133 × 0.5 mm; number of slices = 42 (axial), bandwidth = 50 kHz, GRAPPA acceleration factor = 2. For functional imaging, gradient-echo-based, single-shot echo-planar images covering the whole brain were acquired over multiple sessions (TR = 1500 ms; TE = 15 ms; flip angle = 40°; FOV = 64 × 64 mm; matrix size = 128 × 128; voxel size = 0.5-mm isotropic; number of slices = 42 (axial); bandwidth = 500 kHz; GRAPPA acceleration factor = 2 (anterior-posterior)).

### Analysis

Based on the quality of the images and the eye signal, a total of 10 runs were selected for M1, 10 runs for M2 and 8 runs for M3. Time series were preprocessed using AFNI (Cox, 1996), FSL (Smith et al., 2004), SPM12 (Wellcome Department of Cognitive Neurology, London, United Kingdom) and ANTS software (Advanced Normalization Tools, Avants et al., 2011). For spatial preprocessing, functional volumes were first reoriented, realigned (to correct and estimate motion parameters), resliced and a mean functional image was created for each session using SPM12 (Wellcome Department of Cognitive Neurology, London, United Kingdom). The images were co-registered with the T2-weighted (T2w) structural image (manually skull-stripped) of each individual monkey using the FMRIB’s linear registration tool (FLIRT) of FSL. Functional images were then non-linearly registered to the NIH marmoset brain atlas (Liu et al., 2018) using ANTs. To reduce noise, images were smoothed by a 1.5-mm full-width at half-maximum Gaussian kernel using AFNI. Bold-oxygen-level-dependent (BOLD) response was estimated using a general linear model (GLM) with SPM12 and based on the canonical hemodynamic response reflecting the transient increased blood and nutrient flow following the neuron stimulation.

Fixed-effect individual analyses were performed for each condition in each monkey, with a level of significance set at p < 0.05 corrected for multiple comparisons (FWE, t > 4.8) and at p < 0.001 (uncorrected level, t scores > 3.1). In all analyses, six motions regressors (three rotations and three translations) and regressors of the white matter and cerebrospinal fluid signals were included as covariates of no interest to remove brain motion artifacts and reduce noise. Thanks to the head-fixation system, little to no motion was observed during imaging sessions (see, Schaeffer et al., 2019).

When coordinates are provided in this manuscript, they are presented with respect to the anterior commissure. Results are displayed on coronal sections or fiducial maps obtained with Caret (Van Essen et al., 2001; http://www.nitrc.org/projects/caret/) using the NIH marmoset brain template (Liu et al., 2018).

To define the *encoding looming visual stimuli network*, we contrasted the cortical activations obtained by looming visual stimulations to those obtained by the no-stimuli condition (Figs. 3 and 5). To define the *encoding receding visual stimuli network*, we contrasted the cortical activations obtained by receding visual stimuli to those obtained by the no-stimuli condition (Figs. 4 and 5). To define the *selective looming network*, the above looming visual-stimuli network was additionally masked by the activations obtained for the receding visual-stimuli network (exclusive ‘receding stimulation vs. no stimuli’ mask, p > 0.05 corrected for multiple comparison, FWE, Fig. 6). To define the *selective receding network*, the above receding visual-stimuli network was additionally masked by the activations obtained for the looming visual-stimuli network (exclusive ‘looming stimulation vs. no stimuli’ mask, p > 0.05 corrected for multiple comparison, FWE, Fig. 7).

The quality of the data was assessed by extracting the BOLD time courses from four different regions (Fig. 2). The regions were selected based on the selective looming network contrast performed in group analysis and on the cluster size. The identified regions were the left pulvinar/superior colliculus (27 voxels), the right parietal region (29 voxels), the fundus of the superior temporal sulcus (13 voxels) and the inferior temporal area TE3 (31 voxels).

**Figure 2:**
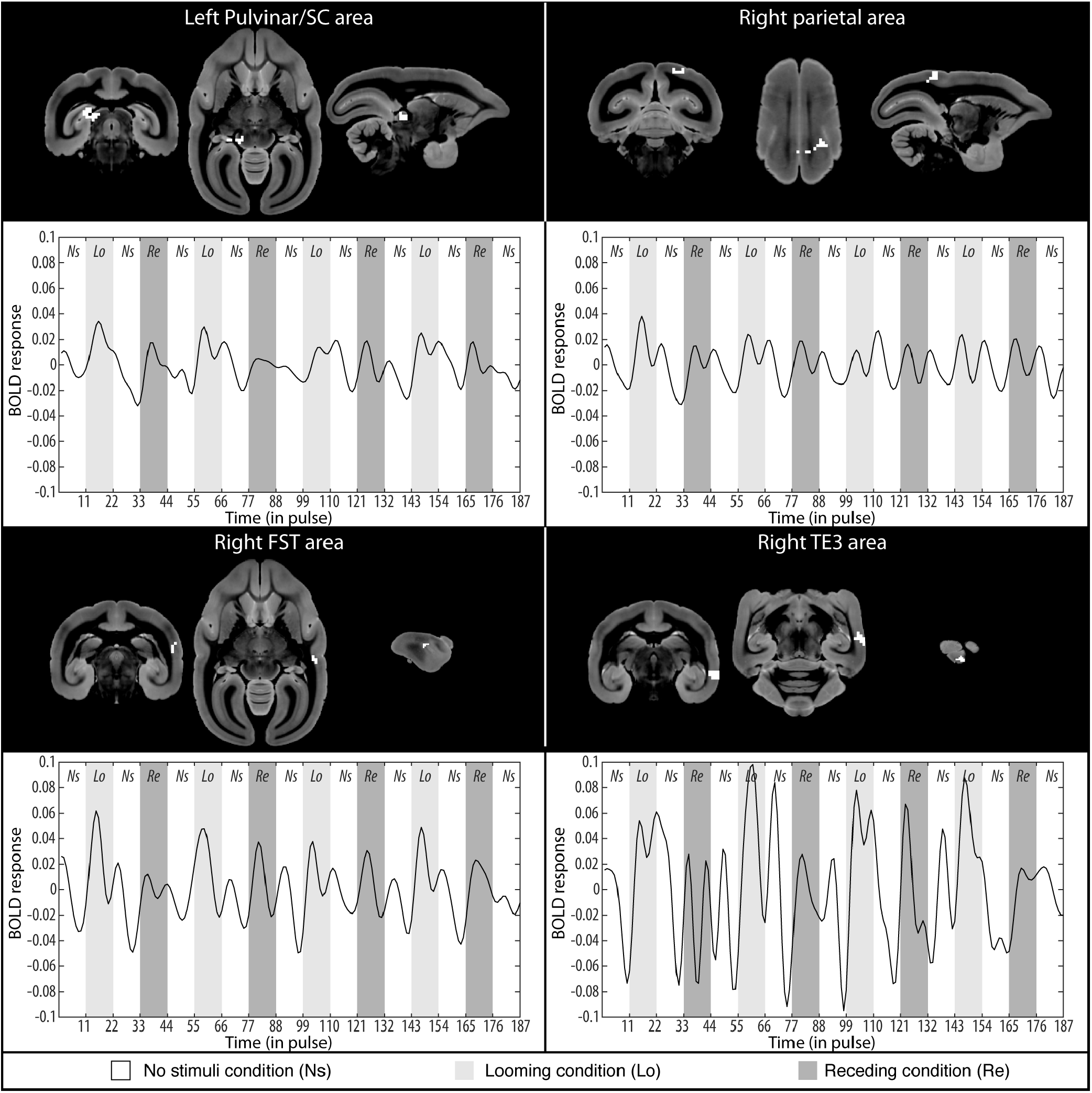
Blood-oxygen-level-dependent (BOLD) time courses extracted in four regions (group average): the left pulvinar/superior colliculus (27 voxels), the right parietal area (29 voxels), the right fundus of the superior temporal sulcus (13 voxels), the right inferior temporal area TE3 (31voxels). For each region, the localisation is shown in coronal, axial and sagittal slices. The light grey shaded area highlights the looming condition block (Lo), the dark grey shaded area the receding condition block (Re) and the no-color area the no-stimulation block (Ns).

The results were consistent across the three animals for the ‘looming vs no stimuli’ and the ‘receding vs. no stimuli’ contrasts, thus Figs. 2, 5, 6, 7 and 8 correspond to a group analysis to simplify the presentation of the results. In the case of Figs 6, 7 and 8, fixed-effect group analyses were performed for each sensory modality with a level of significance set at p < 0.001 (uncorrected level, t scores > 3.1), to observe the maximum activation map (lower power due to the mask constraint) and display on the coronal sections or fiducial map of the NIH marmoset brain template. The labelling refers to the histology-based atlas of Paxinos and colleagues (2011) for the cortical regions (Fig. 1C) and to the atlas of Liu and colleagues (2018) for the subcortical regions.

**Figure 3:**
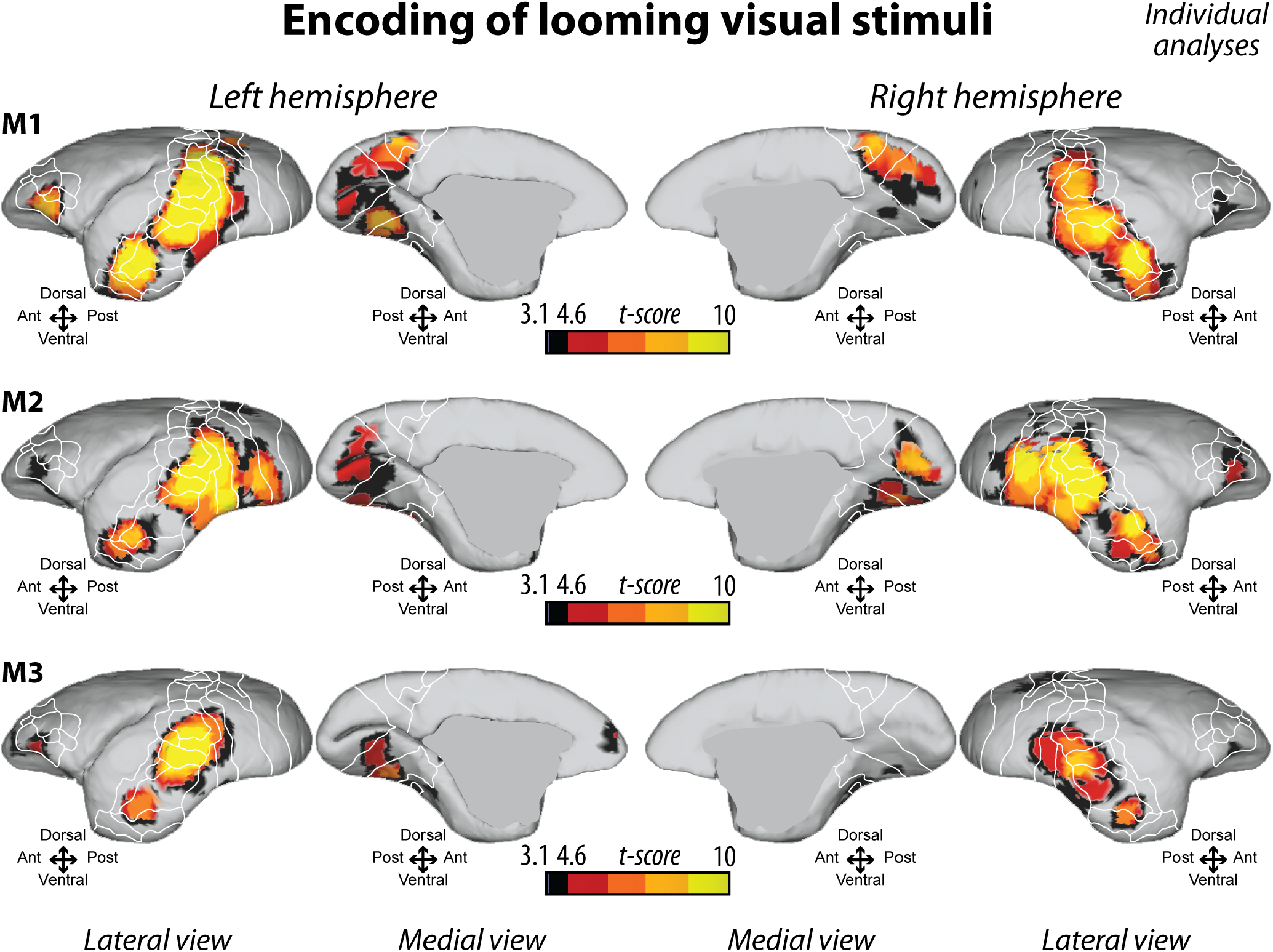
Individual analyses of looming visual-stimuli encoding. For each monkey, activations are warped and presented on the fiducial representation of the NIH marmoset brain template, in the left and the right hemispheres, in medial and lateral views. Black regions correspond to t scores >= 3.1 (p < 0.001 uncorrected level) and t scores < 4.8 (p < 0.05, FWE-corrected level). The regions associated with the red color scale correspond to t scores >= 4.8 (p < 0.05, FWE-corrected level). The white line delineates the regions based on the atlas from Paxinos et al. (2011), see Fig 1C for labelling.

**Figure 4:**
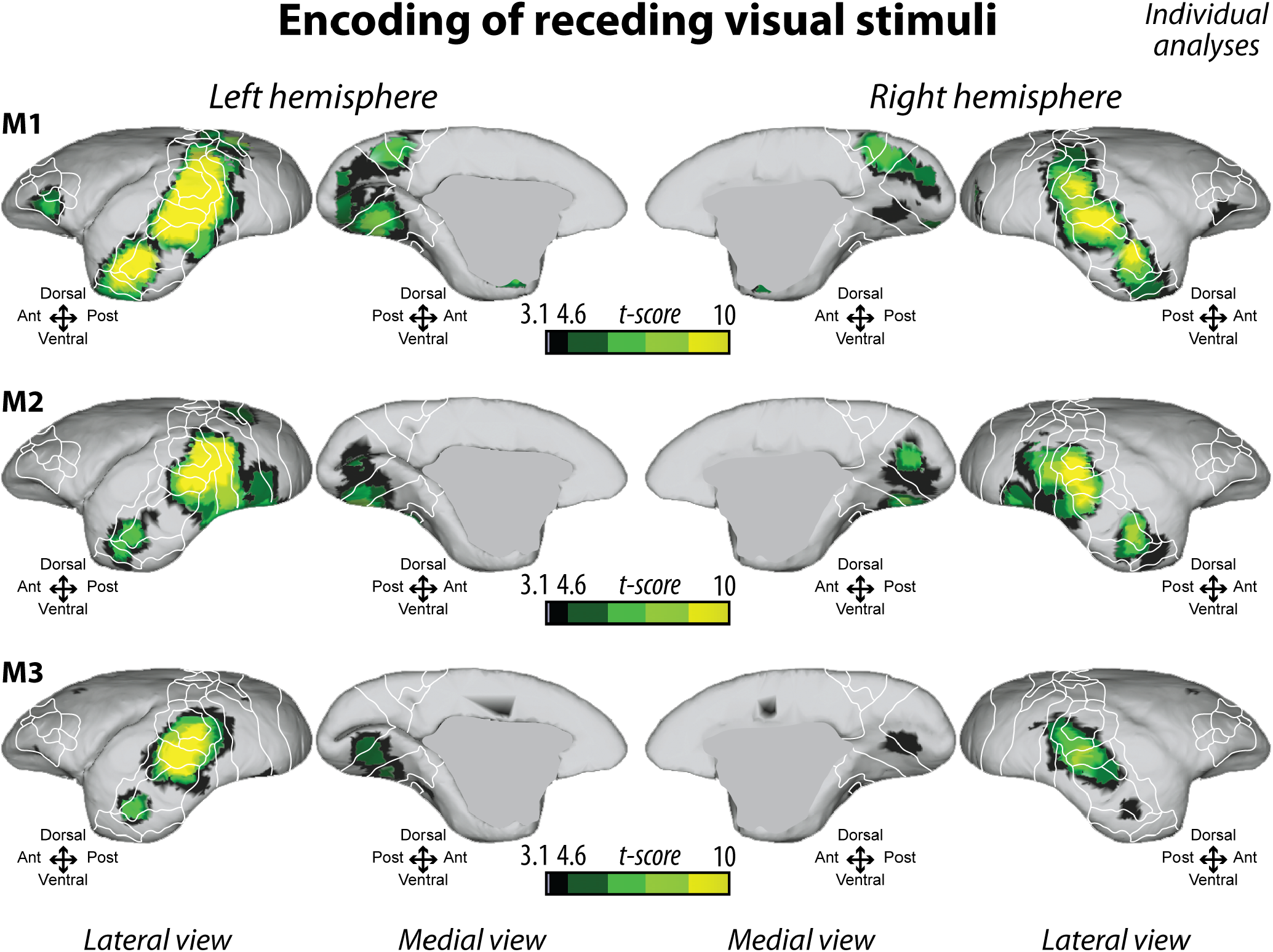
Individual analyses of receding visual-stimuli encoding. For each monkey, activations are warped and presented on the fiducial representation of the NIH marmoset brain template, in the left and the right hemispheres, in medial and lateral views. Black regions correspond to t scores >= 3.1 (p < 0.001 uncorrected level) and t scores < 4.8 (p < 0.05, FWE-corrected level). The regions associated with the green color scale correspond to t scores >= 4.8 (p < 0.05, FWE-corrected level). The white line delineates the regions based on the atlas from Paxinos et al. (2011), see Fig 1C for labelling.

**Figure 5:**
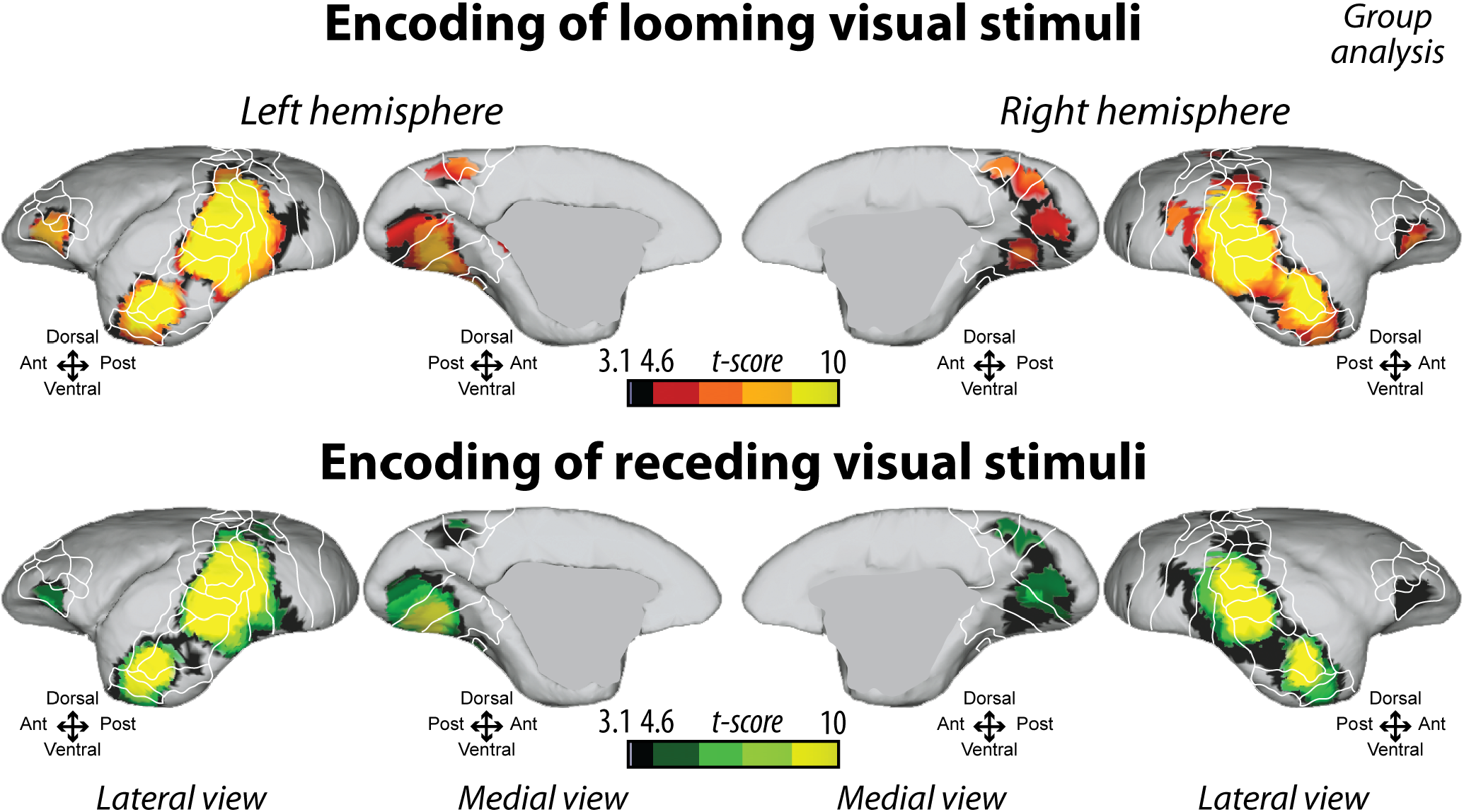
Group analysis of looming (top panel) and receding (low panel) visual-stimuli encoding. For each monkey, activations are warped and presented on the fiducial representation of the NIH marmoset brain template, in the left and the right hemispheres, in medial and lateral views. Black regions correspond to t scores >= 3.1 (p < 0.001 uncorrected level) and t scores < 4.8 (p < 0.05, FWE-corrected level). The regions associated with the red or green color scale correspond to t scores >= 4.8 (p < 0.05, FWE-corrected level). The white line delineates the regions based on the atlas from Paxinos et al. (2011), see Fig 1C for labelling.

**Figure 6:**
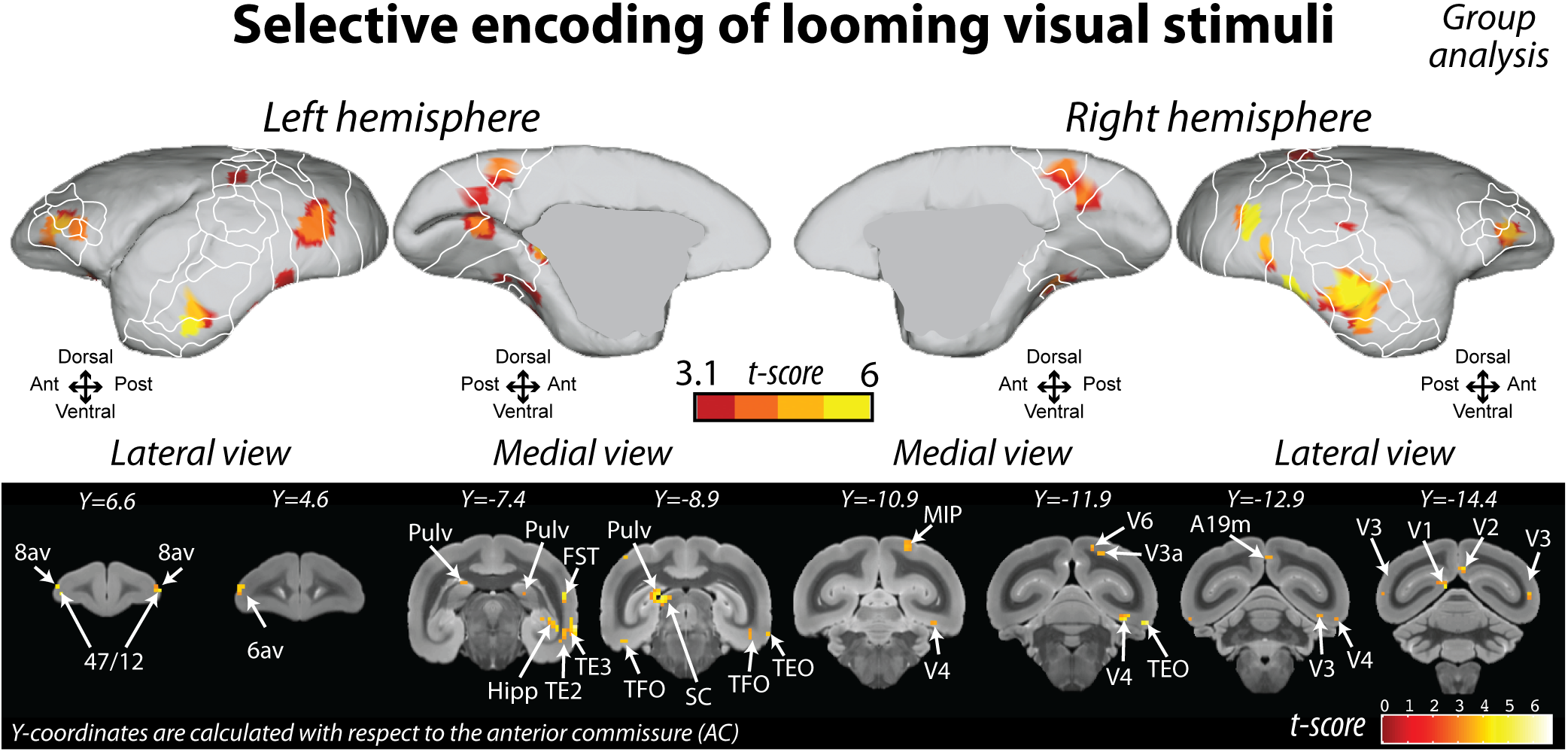
Selective encoding of looming visual stimuli. Upper panel, activations are presented on the fiducial representation of the NIH marmoset brain template (group analysis), in the left and the right hemispheres, in medial and lateral views. The regions associated with the red color scale correspond to t scores >= 3.1 (p < 0.001 uncorrected level, red color scale). Lower panel, activations are presented on the coronal view of the NIH marmoset brain at p < 0.001 uncorrected level. A19m, medial part of area 19; 6Va, area 6 of cortex ventral part a; 8av, ventral part of dorsolateral prefrontal area 8a; 47/12, ventrolateral prefrontal area 47/12; FST, fundus of superior temporal area; Hipp, hippocampus; MIP, medial intraparietal area; Pulv, pulvinar; SC, superior colliculus; TE2, inferior temporal area TE2; TE3, inferior temporal area TE3; TEO, occipital part of temporal area TE; TFO, occipital part of temporal area TF; V1, visual area 1; V2, visual area 2; V3, visual area 3; V3a, visual area 3a; V4, visual area 4; V6, visual area 6. Y-coordinates are calculated with respect to the anterior commissure (in millimeters).

**Figure 7:**
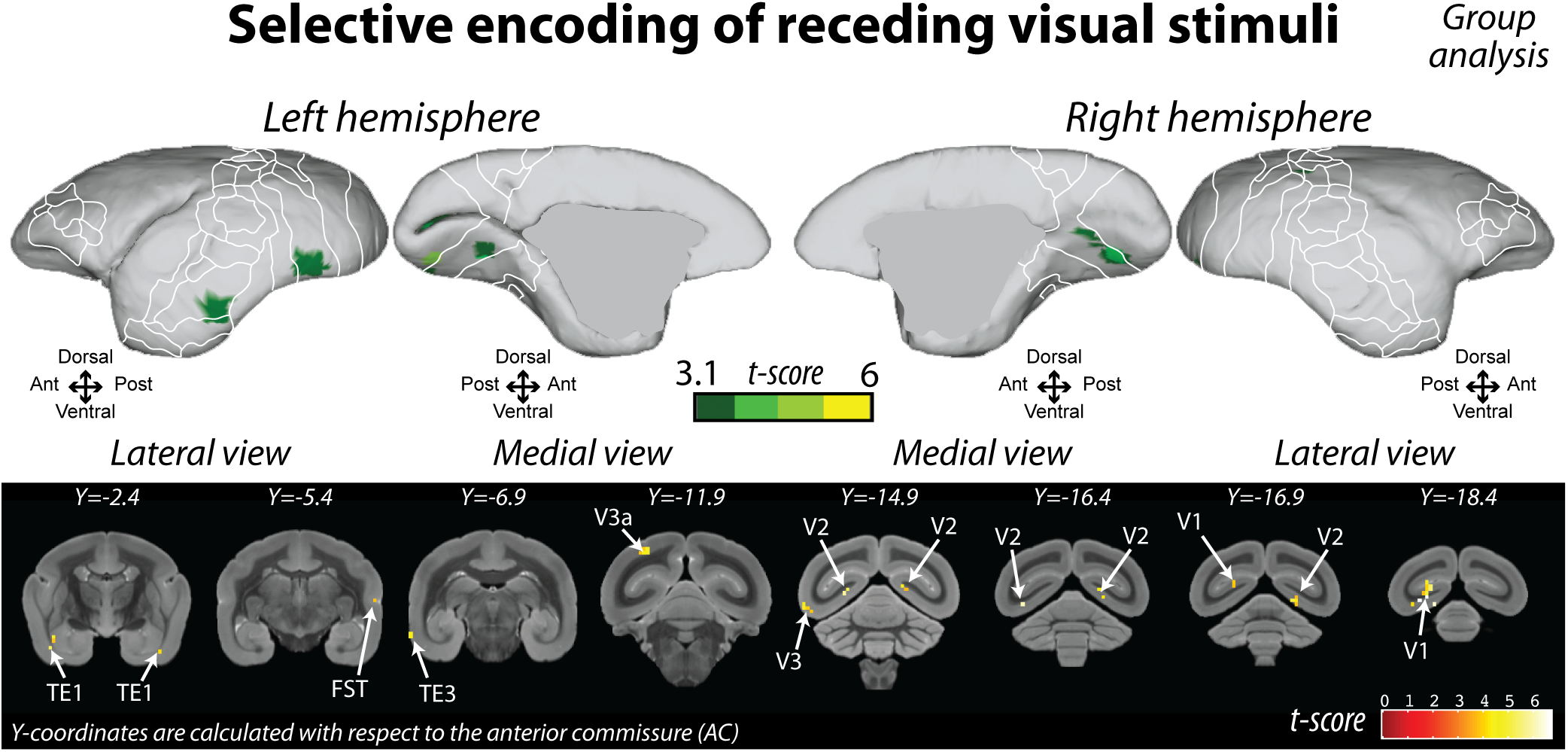
Selective encoding of receding visual stimuli. Upper panel, activations are presented on the fiducial representation of the NIH marmoset brain template (group analysis), in the left and the right hemispheres, in medial and lateral views. The regions associated with the green color scale correspond to t scores >= 3.1 (p < 0.001 uncorrected level, green color scale). Lower panel, activations are presented on the coronal view of the NIH marmoset brain at p < 0.001 uncorrected level. FST, fundus of superior temporal area; TE1, inferior temporal area TE1; TE3, inferior temporal area TE3; V1, visual area 1; V2, visual area 2; V3, visual area 3; V3a, visual area 3a. Y-coordinates are calculated with respect to the anterior commissure (in millimeters).

**Figure 8:**
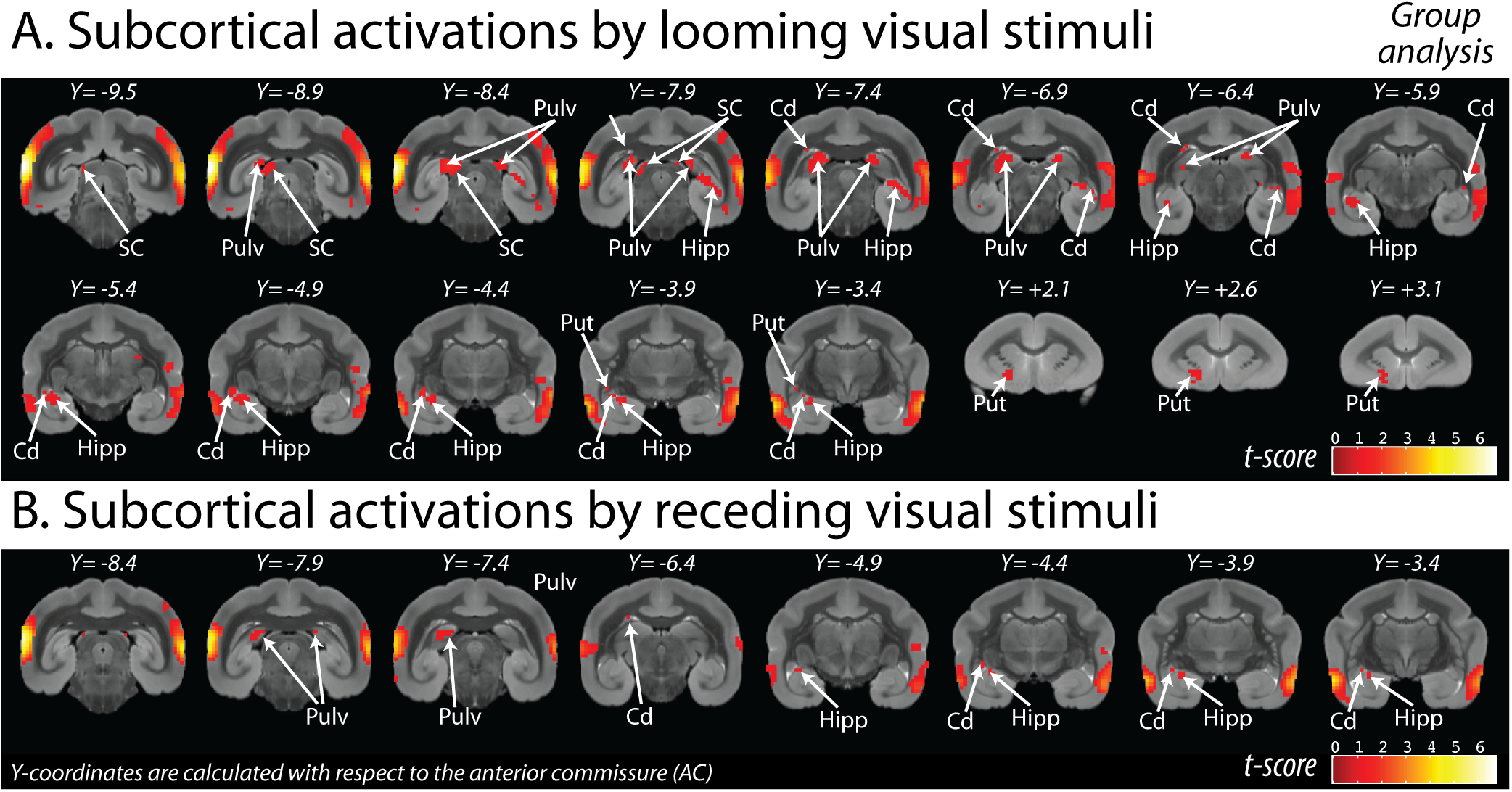
Subcortical regions analysis. A. Activations elicited by visual stimuli looming toward the marmoset. B. Activations elicited by visual stimuli receding away from the marmoset. All activations are presented on the coronal view of the NIH marmoset brain at p < 0.001 uncorrected level (group analysis). Cd, caudate; Hipp, hippocampus; Pulv, pulvinar; Put, putamen; SC, superior colliculus. Y-coordinates are calculated with respect to the anterior commissure (in millimeters).

## Results

In each run, monkeys were presented with three conditions: (1) no visual stimuli blocks; (2) blocks with only visual stimuli looming toward the animal and (3) blocks with only visual stimuli receding away from the animal. In the following, we described the functional cortical and subcortical networks involved in the processing of visual stimuli looming toward or receding away from the subject. The reported activations were identified using an individual analysis for Figs. 3 and 4 and using a group analysis for Fig. 5, both with a level of significance set at p < 0.05, corrected for multiple comparisons (FWE, t-score > 4.8). The reported activations in Figs. 6, 7 and 8 were identified using a group analysis, with a level of significance set at p < 0.001, uncorrected level (t-score > 3.1). As a result, they reflect the activations that are common to the three monkeys involved in the study.

### BOLD response

Figure 2 represents the time courses extracted from four different regions to show how the BOLD response changed during the task. The regions of interest were defined from the selective looming contrast obtained by the group dataset. In the four selected regions, the BOLD response increased during the visual looming condition (Fig. 2, light grey shaded area) and decreased during the no-stimuli condition. During the visual receding condition (Fig. 2, dark grey shaded area), some increases of the BOLD response were observed, but with a lower amplitude compared to the looming condition. The second jump observed at the beginning of the no-stimuli condition in FST and parietal areas is probably related to the reward delivery (between each block).

### Looming visual stimulations

Visual stimuli looming toward the animal (‘Visual looming vs. no stimuli’ contrast) activated a large extent of the temporal cortex and the occipital cortex as well as some portions of the parietal cortex and the prefrontal cortex (Fig. 3, individual analyses, Fig. 5, group analysis). By masking this contrast with the activations obtained during the receding conditions (‘Visual looming vs. no stimuli’ masked exclusively by ‘Visual receding vs. no stimuli), we revealed the *selective visual looming network* (Fig. 6, upper part, fiducial view, group analysis). This means that these voxels were activated by visual looming stimuli and not by receding stimuli. More specifically, this network consists of some portions of the temporal cortex (fundus of superior temporal area FST; inferior temporal areas TE2, TE3; occipital part of temporal areas TE and TF), occipital cortex (visual areas V1, V2, V3, V3a and V4; dorsomedial area V6 and medial part of area 19), prefrontal cortex (ventrolateral prefrontal area 47/12 and 45, ventral part of areas 8a and 6a) and medial intraparietal area MIP (Fig. 6, lower part, coronal slices). We also observed subcortical activations in the superior colliculus (SC), the pulvinar (Pulv) and the hippocampus (Hipp).

### Receding visual stimulations

Visual stimuli receding away from the animal (‘Visual receding vs. no stimuli’ contrast) activated a large extent of the temporal cortex and the occipital cortex as well as some parietal areas (Fig. 4, individual analyses, Fig. 5, group analysis). By masking this contrast with the activations obtained during receding conditions (‘Visual receding vs. no stimuli’ masked exclusively by ‘Visual looming vs. no stimuli’), we revealed the *selective visual receding network* (Fig. 7, upper part, fiducial view, group analysis). These voxels were activated by visual receding stimulation and not by looming stimulations. More specifically, this network consists of some portions of the lateral temporal cortex (fundus of superior temporal area FST), inferior temporal cortex (TE1 and TE3), and occipital cortex (visual area V1, V3, V3, V3a) (Fig. 7, lower part, coronal slices).

### Subcortical activations

The high resolution of the imaging acquisition allowed us to capture subcortical activations evoked by visual looming (Fig. 8A) or receding stimuli (Fig. 8B). Both types of visual stimuli elicited activations in the caudate, the hippocampus and the pulvinar. In most cases, the subcortical activations were stronger and more widespread in the looming compared to the receding condition. Visual looming stimuli also triggered activations in the putamen and the superior colliculus.

## Discussion

Here we explored the neural activations elicited by visual stimuli looming toward or receding away from awake marmosets using fMRI at 9.4T. Both visual stimuli activated a large cortical network in frontal, parietal, temporal and occipital cortex. The network evoked by looming stimuli was stronger and showed some selective encoding of subcortical areas, including the pulvinar and the superior colliculus.

### Ventral visual pathways for object features

We found robust activations in the occipitotemporal cortex including V1, V2, V3, V4, TEO and TE for looming and receding stimuli. This network is very similar to the one found by Hung and colleagues (2015a) that investigated the responses to static visual stimuli (faces, body, objects) with fMRI in marmosets. Areas V4 and TEO both showed preferential encoding for complex visual stimuli (Hung et al., 2015a, 2015b). Moreover, lesions in the marmoset inferior temporal cortex indicate its involvement in visual object discrimination (Ridley et al., 2001). Thus, when visual stimuli loom toward the monkey or recede away from it, activation of these visual areas likely reflects object analysis.

### Dorsal visual pathways for motion selectivity

Interestingly, in the fMRI study of Hung and colleagues (2015a), no activations were found in medial and superior temporal areas or in parietal cortex. However, in our current study, in addition to the visual areas V1, V2 and V3, we observed activations in areas MT, MST, FST and parietal cortex (Opt). These areas compose the dorsal visual pathway involved in the guidance of visually directed behavior and spatial orienting (Goodale and Milner, 1992). Some V1 neurons are speed sensitive (Yu et al., 2010), some are motion sensitive and depend on contour orientation extraction, but others are independent and seem involved in complex motion analysis (Tinsley et al., 2003; Barraclough et al., 2006; Guo et al., 2006). V2 neurons showed orientation and direction selectivity (Lui et al., 2005; Barraclough et al., 2006). V3 neurons mainly encode central vision and are orientation selective but direction insensitive (Rosa and Tweedale, 2000). Area MT seems to strongly resemble its macaque homolog and plays a crucial role in motion analysis. MT neurons show direction selectivity, contrast sensitivity and spatial frequency preferences (Rosa and Elston, 1998; Solomon et al., 2011; Liu et al., 2013). Areas MST and FST also show motion and direction selectivity (Krubitzer and Kaas, 1990; Palmer and Rosa, 2006a, 2006b) and have been suggested to play a critical role in higher stages of motion processing by sending strong feedback to area MT and other cortical areas (Palmer and Rosa, 2006a; Solomon and Rosa, 2014).

The fact that Hung and colleagues (2015a) did not observe these activations is probably due to the use of static images, whereas here, our dynamic visual stimuli activated motion-sensitive areas. Interestingly, both looming and receding stimuli seem to activate the ventral and dorsal visual pathways. Over the past years, more evidence of strong interconnections between the two visual pathways has accumulated (for review, see Handa and Mikami, 2018). The authors suggest that these interconnections allow sharing of information about motion and shape between the early visual areas in both pathways with a mechanism relying on visual cues and behavioral requirements.

### Looming selective network

We observed some areas that showed a preference for encoding looming visual stimuli. These included some parts of ventral visual areas, V4, TE, TPO involved in visual object recognition, properties and discrimination as shown in marmosets (Ridley et al., 2001; Hung et al., 2015a), macaques (Desimone and Schein, 1987; Schein and Desimone, 1990; Okazawa et al., 2015; El-Shamayleh and Pasupathy, 2016) and humans (Winawer and Witthoft, 2015). This ventral stream seems essential for quickly extracting these features of the incoming stimuli and for determining whether they pose a potential threat.

We also observed selective activation in frontal areas 6Va and 8av. The marmoset area 6Va (Burman et al., 2006; Paxinos et al., 2011) is similar to its homolog in macaques (Barbas and Pandya, 1987) also called F4 (Matelli et al., 1985). In marmosets, stimulation in this area led to some facial and forelimb movements (Burman et al., 2008). This suggests that stimuli looming toward the marmoset can elicit facial expression and participate in the planning of future movement. The marmoset area 8av has been suggested to be the homologous of the macaque frontal eye field (FEF) with strong anatomical connections to the dorsal stream, especially MT, MIP, V6, extrastriate visual areas (Burman et al., 2006; Reser et al., 2013) and functional connections to the areas 6Va, 45 and V4 (Ghahremani et al., 2017). This connectivity suggests an involvement in spatial cognition, attention and eye movements.

We also observed selective activation in parietal area MIP and area V6. In macaques, MIP neurons respond to the spatial parameters of planned goal-directed movements (Kuang et al., 2016), including the physical aspect of the planned movement (e.g. motor preparation) and the visual aspect of the planned movement (e.g. upcoming movement kinematics). Area V6 is involved in visual motion and action both in macaques (Galletti et al., 1999; Fan et al., 2015) and in humans (Pitzalis et al., 2013a, 2013b, 2013c). It has been suggested that V6 is involved in processing visual motion signals and conveying to V6A the visual and oculomotor information to orchestrate the eye/arm movements required to avoid or reach moving or static objects (Galletti et al., 1999; Fan et al., 2015; Pitzalis et al., 2015). In macaques, both areas MIP and V6 receive strong inputs from visual areas. MIP is strongly connected to V6 and to the dorsal premotor cortex (Galletti et al., 2001; Bakola et al., 2017). In marmosets, the area V6 receives input mainly from visual areas, MT and MIP (Rosa et al., 2009). Little is known about the marmoset parietal cortex; however, the dorsal premotor cortex receives very sparse inputs from MIP (Burman et al., 2014), pointing to a potential difference in cortical organization between Old and New World monkeys. The strong connection of MIP and V6 to the middle temporal area in marmosets (Abe et al., 2018) suggests that, in this species, these areas are involved in visual motion analysis. Interestingly, an fMRI study in humans showed the involvement of V6 for visual stimuli looming toward the near space of the subject (Quinlan and Culham, 2007).

The network activated by visual stimuli looming toward the marmoset showed similarities with the macaque network involved in naturalistic 3D visual looming objects (Cléry et al., 2018) and in visuo-tactile impact prediction (Cléry et al., 2017; Cléry and Ben Hamed, 2018) with the activations of MIP, 12, FEF (8av), F4 (6Va), FST, TE, TPO, and visual areas. These similarities suggest that a part of the alert network involved in impact prediction and peripersonal space encoding is also involved in the processing of visual looming stimuli.

### Subcortical areas

Both in rodents (Sahibzada et al., 1986) and even in macaques (DesJardin et al., 2013), SC stimulation can induce defensive-like behaviors. In rodents, the SC has looming-sensitive neurons (Westby et al., 1990) and coordinates this defensive behavior by transferring threat signal to the parabigeminal nucleus or the lateral posterior thalamic nucleus (LPTN) to activate a fleeing or freezing response (Shang et al., 2018). In marmosets, SC neurons show direction selectivity and have been suggested to play a role in the processing of looming stimuli (Tailby et al., 2012). The SC is strongly connected to the pulvinar (Stepniewska et al., 2000; Ghahremani et al., 2017; Kwan et al., 2019), the primate homologous of LPTN, where we also observed activations, suggesting a freezing behavior, consistent with the fact that the animal is restrained and cannot escape. These findings suggest a potentially preserved subcortical networking underlying freezing responses in rodents and primates.

### Receding selective network

We observed few activations selective to the receding stimuli and most were confined to the inferior temporal cortex (TE) and occipital cortex (V1, V2, V3). Similarly, the receding of a 3D object from the near space elicited few activations, mostly in the temporal cortex, in macaques (Cléry et al., 2018). This suggests that the majority of the regions activated by receding stimuli are also activated by looming stimuli and reflects the importance of the information brought by the direction of moving stimuli.

In summary, visual stimuli looming towards marmosets activated a large cortical and subcortical network composed of the pulvinar, superior colliculus, prefrontal cortex and temporal cortical areas. The strong connections between these areas and their functions suggest the existence of an alert network processing the visual stimuli looming toward our peripersonal space, to extract the motion, orientation and identity of the visual stimuli to evaluate the potential consequences of the moving stimuli (impact, avoidance, grasping). Depending on the context, this alert network prepares the body for protective behaviors (e.g. fleeing or freezing) and emotional components (e.g. anxiety, fear). This network seems to be conserved across New and Old-World primate species and supports the view that marmosets are a viable model to study visual and multisensory processes by using fMRI to guide further invasive recordings and/or pharmacological manipulations.

## Acknowledgements

Support was provided by the Canadian Institutes of Health Research (FRN 148365) and the Canada First Research Excellence Fund to BrainsCAN. We thank Drs. Cirong Liu and Afonso Silva of National Institutes of Health for providing the surface models of the marmoset brain. We also thank Miranda Bellyou for animal preparation and care and Dr. Alex Li for scanning assistance.

